# INHBA promotes the progression of gastric cancer by activating MAPK signaling pathway via targeting ITGA6

**DOI:** 10.1101/2025.04.05.647266

**Authors:** Guojian Zhou, Rui Zhang, Lei Nie, Yi Si, Ting Liu, Jing Wang, Shuangshuang Han, Mingda Xuan, Weifang Yu, Jia Wang

## Abstract

Gastric cancer (GC) is one of the most common malignancies, ranking as the fifth most common cancer and the fourth leading cause among cancer related deaths worldwide. Upregulated INHBA expression in gastric cancer tissues compared to adjacent non-cancerous tissues was confirmed by immunohistochemistry and qRT-PCR analysis. This increased expression of INHBA was found to be significantly associated with the incidence of tumor lesion, lymph node metastasis and the progression to more advanced TNM stages in patients. Functional experiments showed that INHBA could promote the proliferation of GC cells, enhance migration and invasion *in vitro*, while simultaneously facilitating the inhibition of apoptosis. To test whether INHBA regulated the tumor growth *in vivo*, the animal studies were performed. The indicated that INHBA over-expression promoted the tumor growth, including weight and volume. Moreover, this study demonstrated through a series of experiments including RNA-seq, Co-IP, Co-IF, Western blot, and Rescue studies that INHBA promotes the progression of GC by targeting ITGA6 to regulate the MAPK signaling pathway. In summary, understanding the role of INHBA/ITGA6/MAPK in tumourigenesis could provide new insights into gastric cancer therapy and targeted inhibition of INHBA might be a potential therapeutic approach for GC treatment.

## Introduction

Gastric cancer remains a significant global health burden, representing one of the most prevalent malignancies worldwide. According to recent GLOBOCAN estimates, it ranks as the fifth most commonly diagnosed cancer and the fifth leading cause of cancer-related mortality globally, accounting for approximately 1 in 20 new cancer cases and 1 in 16 cancer deaths annually[1]. The pathogenesis and progression of GC are mediated through a complex interplay of molecular mechanisms, gene-related factors, and environmental influences.Individuals diagnosed with late-stage GC face dismal clinical outcomes, typically exhibiting a median overall survival (OS) under 12 months[2]. Therefore, understanding how gastric cancer develops and finding new treatment gene-targets are critical to improving patient care.

Inhibin beta A (INHBA), a member of the transforming growth factor-beta (TGF-β) superfamily, functions as a heterodimeric protein composed of α and βA subunits. Identified in 1978 as a key modulator of the hypothalamic–pituitary–gonadal(HPG) axis, INHBA forms a disulfide-linked homodimer called activin A[3]. Several studies found that abnormal expression of INHBA exerted various biological functions in different tumor development processes including colorectal cancer, esophageal carcinoma, prostate, breast, Cervical and ovarian cancer[4–9].Studies have revealed that INHBA expression correlates with cancer progression and poor prognosis. Emerged mechanistic studies suggested INHBA’s role in oncogenic processing may involve epithelial-mesenchymal transition (EMT) through typical TGF-β/Smad signaling axis[10–11]. The potential of INHBA as a therapeutic target for cancer treatment has also been validated, with some promising results in early-phase studies[12].Therefore, these results indicated that INHBA is related to the occurrence, development and prognosis of various cancers.

At present, there are few studies on INHBA in gastric cancer. Elevated INHBA expression correlated with reduced 5-year survival rates, demonstrating biomarker potential in gastric adenocarcinoma[13–14]. Several studies indicated that some circular RNA and transcription factors can regulate the expression of INHBA in different ways, thus affecting the biological function of gastric cancer cells and driving the progression of GC[15–17]. Research have shown that INHBA gene inhibited the migration and invasion of gastric cancer cells by blocking the activation of TGF-β signaling pathway[18]. Emerging evidence positions INHBA as a potential modulator in gastric cancer, yet its precise oncogenic signatures in gastric cancer persist as an underdeveloped research frontier, with only TGF-β pathway studies addressing this target in GC models.

Therefore, This study elucidated INHBA-driven oncogenesis via one type of signaling pathway and downstream gene-effector regulation.We found that INHBA is highly expressed in gastric cancer and is correlated with poor survival prognosis in gastric cancer patients. INHBA enhances the malignant phenotype of gastric cancer cells and promotes the growth of gastric cancer *in vivo*. Integrative analysis of transcriptome sequencing, bioinformatics analysis, Co-IP/Co-IF experiments, and Rescue studies, we first identified ITGA6 as the dominant INHBA effector in gastric cancer. We also demonstrated ITGA6 is highly expressed in gastric cancer and interacts with INHBA. INHBA upregulates the expression of ITGA6 and activates the typical MAPK signaling pathway, further promoting the proliferation, migration, and invasion capacities of gastric cancer cells, ultimately accelerating the progression of gastric cancer.

## Materials and Methods

### Patients and samples

A total of 50 paired GC and adjacent normal tissues were surgically resected from patients at the First Hospital of Hebei Medical University. No patients received chemotherapy/radiation before surgery, and all provided informed consent through hospital ethics protocols. Isolated specimens were snap-frozen in liquid nitrogen and stored at Hebei Medical University First Hospital Biobank. Histologically confirmed gastric adenocarcinoma (including signet-ring cell carcinoma subtypes) were used for tumor tissue analysis in this study. The study protocol received ethical approval (Approval No.S00996)) from Hebei Medical University First Hospital Ethics Committee in accordance with the Declaration of Helsinki.

### Cell culture

Shanghai iCell Bioscience Inc provided the GC cells (NUGC-3, Hs746T) and Wuhan Pricella Biotechnology Co., Ltd. provided the human gastric mucosa cell (GES-1) and GC cells (HGC-27, AGS), which were confirmed by STR analysis. The cells were maintained in RPMI 1640 or DMEM medium (Gibco, Gaithersburg, MD, USA) supplemented with 10% fetal bovine serum (FBS; Gibco) and 1% penicillin-streptomycin solution (Solarbio Science & Technology Co., Ltd., Beijing, China). Cells were cultured in a 37°C incubator with 5% CO₂.Monthly mycoplasma screening was performed by PCR, while cell passage numbers was strictly controlled within 30 passages.

### Transfections, stable cell lines, siRNAs, shRNAs, and plasmids

Transfections were performed with Lipofectamine 3000 (Invitrogen; Carlsbad, CA, USA) when cells reached 60–70% confluence. NUGC-3/Hs746T and HGC-27/AGS cells were respectively transfected with INHBA-targeting siRNA or overexpression plasmids (with negative controls; GenePharma Co., Ltd., Shanghai, China). AGS cells were transfected with siRNA-ITGA6 and siRNA-NC, while NUGC-3 cells were transfected with a ITGA6 overexpression plasmid and GenePharma negative control. Functional assays were conducted 48h post-transfection. For stable clone selection, neomycin (600 μg/mL) was introduced into HGC-27 cultures 48h after INHBA-overexpressing plasmid transfection, generating oe-Vector and oe-INHBA isogenic cell lines (antibiotic-free background). Upon monoclonal colony establishment, cells were maintained in 300μg/mL G418-supplemented medium for 10 consecutive days to stabilize transgenic expression.

### RNA extraction and qRT-PCR

Total RNA was isolated from cellular and tissue samples using RNA-Easy Isolation Reagent (Vazyme Biotech Co., Ltd., Nanjing, China). Reverse transcription was performed with the PrimeScript RT Reagent Kit (Takara Bio, Beijing, China) to synthesize complementary DNA (cDNA). For quantitative real-time PCR (qRT-PCR), reactions were carried out using the AceQ Universal SYBR qPCR Master Mix (Vazyme). β-actin mRNA served as the endogenous reference for normalization. All experiments included three technical replicates per sample. The primers used are shown in Supplementary Table 1. Cycle threshold (CT) values were analyzed using the 2^-ΔΔCT method to calculate relative mRNA expression levels.

### Immunohistochemistry (IHC)

Tissue samples from patients and nude mice were fixed in 4% paraformaldehyde (PFA) solution, followed by paraffin embedding using standard histological procedures (Shanghai YiYang Instrument Co., Ltd., China). Subsequently, 4-μm thick sections were prepared and subjected to staining. For immunohistochemical analysis, sections were processed using the Rabbit two-step detection kit (ZSGB-Bio, Beijing, China) according to the manufacturer’s protocol. The following primary antibodies from Proteintech (Wuhan, China) were employed: INHBA (diluted 1:200; Proteintech; Catalog number: 17524-1-AP), Ki67 (diluted 1:100; Proteintech;Catalog number: 27309-1-AP) and ITGA6 (diluted 1:500; Proteintech ; Catalog number: 27189-1-AP). Staining results were quantitatively analyzed using the average optical density (AOD) method combined with a double-blind scoring system.

### Western blot analysis and antibodies

Protein extraction was performed using RIPA lysis buffer (Solarbio) mixed with protease/phosphatase inhibitor (Solarbio) at a 100:1 ratio. Proteins were separated by electrophoresis on a 10% SDS-polyacrylamide gel (Bio-Rad Laboratories Inc., Hercules, CA, USA) and subsequently transferred to a polyvinylidene fluoride (PVDF) membrane (Merck Millipore, Billerica, MA, USA). In this study, the following antibodies were used: GAPDH antibody (diluted 1:500; Goodhere-Bio; Catalog number: AB-P-R001), INHBA antibody (diluted 1:500; Proteintech, Wuhan, Hubei, P.R.C; Catalog number: 17524-1-AP), ITGA6 antibody (diluted 1:500; Proteintech, Catalog number: 27189-1-AP), MEK1/2 antibody (diluted 1:1000; Proteintech; Catalog number: 11049-1-AP), phospho-MEK1 antibody (diluted 1:1000; Proteintech; Catalog number: 28930-1-AP), ERK1/2 antibody (diluted 1:1000; Proteintech; Catalog number: 66192-1-Ig). phospho-ERK1/2 antibody (diluted 1:100; Proteintech; Catalog number: 28733-1-AP). The Odyssey Scanning System (LI-COR Biosciences, Lincoln, NE, USA) identified the immunoreactive protein bands.

### Cell proliferation assay

For the CCK-8 assay, cells from each group were seeded evenly in 96-well plates at a density of 2 × 10³ cells per well and cultured in complete medium. At 24, 48, 72, and 96 hours post-seeding, 10 μL of Cell Counting Kit -8 (CCK-8; Dojindo, Tokyo, Japan) reagent was added to each well. After an additional 2-hour incubation, the absorbance at 450 nm was measured using a Promega GloMax luminescence detector (Promega, Madison, WI, USA).

For the colony formation assay, cells were plated at a density of 1000 cells per well in 6-well plates and cultured in complete medium, which was replaced every three days. After seven to fourteen days, the cells were washed twice with PBS, fixed with 4% paraformaldehyde for 30 minutes, and stained with 0.1% crystal violet for 20 minutes.

### Cell migration assay

For the wound healing assay, cells from each group were evenly seeded in 6-well plates. When cell confluence reached approximately 100%, a 200 μL pipette tip was used to drow two straight lines in each well to simulate wounds. After washing twice with PBS, the cells were cultured in serum-free medium. Wound images were captured at 0 and 48 hours, and the migration rate was calculated as the ratio of the gap width at 0 hours to that at 48 hours.

For the Transwell migration assay, 4×10^4^ (HGC-27, AGS) or 6×10^4^ (NUGC-3, Hs746T) cells in 200 μL of serum-free medium were added to the upper chamber (Corning Incorporated, Corning, NY, USA), while 700 μL of complete medium was added to the lower chamber to induce cell migration. After 48 hours of incubation, cells on the upper side of the polycarbonate membrane were removed with a cotton swab. The membrane was washed twice with PBS, fixed with 4% paraformaldehyde for 30 minutes, and stained with 0.1% crystal violet for 20 minutes. Excess stain was removed by washing with PBS. Five randomly selected fields were photographed, and the stained cells were counted using ImageJ software (National Institutes of Health, Bethesda, MD, USA).

### Cell invasion assay

To facilitate downward invasion, gastric cancer cells were resuspended in 100 μL of serum-free medium and seeded into the upper chamber, while 600 μL of complete medium was added to the lower chamber for the Transwell invasion assay. All other experimental procedures followed the same protocol as the Transwell migration assay.

### Cell apoptosis assay

After 48 hours of transfection, cells were collected from each group and stained with the Annexin V-FITC/PI apoptosis assay kit (NeoBioscience, Shenzhen, China). A total of 1×10^5^ cells were analyzed for apoptosis by flow cytometry ( BD Biosciences, San Jose, CA, USA) and the apoptotic rate was calculated using FlowJo software.

### Animal studies

Ten male BALB/c nude mice, weighing between 12-16g and aged 4-5 weeks, were acquired from Beijing Sibeifu Biotechnology Co., Ltd. They were randomly divided into two groups, each consisting of five animals, and housed in a pathogen-free environment with free access to food and water. The temperature was maintained at approximately 22°C, with a 12-hour light/dark cycle.After disinfecting the skin of the nude mice’s left hind limb flank, these nude mice were subcutaneously inoculated with stable overexprssion of INHBA and negative control HGC-27 cells (6×10^6^ cells per mouse) by two sterile syringe. Beginning on the fourth day after injection, tumor size were measured every two days, and the volume was determined using the formula volume = (long diameter × short diameter²)/2. The mice were humanely euthanized when the tumor volume approached 1000 mm³. The tumor tissues were then sectioned, embedded in paraffin, and subjected to HE staining and IHC examination. The study protocols were approved by the Experimental Animal Care and Use Committee and the Ethics Committee of Hebei Medical University.

### RNA sequencing (RNA-seq)

Following 24-hour transfection with siRNA-NC and siRNA-INHBA, total RNA was extracted from NUGC-3 cells. RNA library preparation and sequencing analysis were performed by Beijing Novogene Technology Co, Ltd. Briefly, six samples (triplicates of siNC and siINHBA) were sequenced on the Illumina NovaSeq 6000 platform. Raw sequencing data underwent quality control and filtering to obtain high-quality reads, which were then aligned to reference genomes using HISAT2 software for gene expression quantification. Differentially expressed genes (DEGs) were identified using DESeq2 software with the threshold criteria of log2 |FoldChange| > 1.0 and P-value < 0.05.

### Co-Immunofluorescent (Co-IF) assay

NUGC-3 cells were evenly seeded on coverslips in a 6-well plate. At 20-30% confluence, cells were gently washed three times with PBS. Following PBS washes, cells were sequentially processed through the following steps: blocked with 2% bovine serum albumin (BSA-V;Solarbio; Catalog number: A8020), fixed with 4% paraformaldehyde and penetrated with 0.2% Triton X-10 (Solarbio; Catalog number: T8200). Primary INHBA antibody (diluted 1:50; Proteintech; Catalog number: 17524-1-AP) and ITGA6 antibody (diluted 1:50; Santa Cruz Biotechnology; Catalog number: sc-374057) were added to the cells and incubated overnight at 4°C. After primary antibody incubation, cells were incubated with fluorescently-labeled secondary antibodies (1:500 dilution; Cy3 anti-rabbit IgG, FITC anti-mouse IgG; Beyotime Biotechnology Co.Shanghai, China) for an hour at room temperature in the dark condition. Finally, nuclei were stained with 4’6’-diamino-2-phenylindole dihydrochloride (DAPI; Beyotime Biotechnology Co.). Then the representative images were acquired using a fluorescence microscope.

### Co-immunoprecipitation (Co-IP)

NUGC-3 cells were washed twice with PBS using gentle condition. Cell lysis was performed according to the manufacturer’s protocol using the Pierce™ Classic Magnetic IP/Co-IP Kit (Thermo Scientific, Waltham, MA, USA; Cat# 88804). After cell lysis, the lysate was mixed with 10 μg of INHBA antibody, ITGA6 antibody and IgG control antibody from Proteintech. The mixture was subsequently incubated at room temperature for two hours with gentle rotation. Following the addition of magnetic beads to the lysate, the mixture was continuously rotated at room temperature for an hours. Proteins were eluted using Elution Buffer following magnetic bead separation. The pH was neutralized with Neutralization Buffer. The resulting samples were analyzed by Western blot analysis.

### Bioinformatics analysis

The expression levels of INHBA mRNA in gastric cancer tissues and adjacent normal tissues were analyzed using the GEO (https://www.aclbi.com/static/index.html#/geo) database. Gene expression levels and survival analysis in GC were analyzed using GEPIA2 (http://gepia2.cancer-pku.cn/#index), while subcellular localization was analyzed by GeneCards (https://www.genecards.org/). The specific primers were designed and validated using PrimerBank (https://pga.mgh. harvard.edu/primerbank/) in combination with an advanced primer design tool (https://www.nc\bi.nlm.nih.gov/tools/primer-blast/). KEGG enrichmentt analysis were performed on the Database for Annotation, Visualization, and Integrated Discovery (David database).

### Statistical analysis

Three independent experimental replicates were performed for each group to ensure the reliability of results, in accordance with established guidelines in the field. Data following a normal distribution were expressed mean ± standard, whereas non-normal data utilized median and interquartile distance. SPSS 26.0 (IBM, Armonk, NY, USA) and GraphPad Prism 9.5 (GraphPad Software, La Jolla, CA, USA) were used for statistical analysis in this study. The statistical analyses were performed using Student’s *t*-test for pairwise comparisons, one-way ANOVA for single-factor group comparisons, and two-way ANOVA for assessing interactions between two independent variables. *P* < 0.05 was considered statistically significant.

### Data Availability

The data that supporting the findings of this study are available from the corresponding author upon request. The Gene Expression Profiling Interactive Analysis (GEPIA2) and Gene Expression Omnibus (GEO) at GSE63089 and GSE66229 provided the data used in this analysis.

## Results

### INHBA is upregulated in GC patients and is associated with poor prognosis

To elucidate the role of INHBA in the pathophysiology of gastric GC, we initially examined its expression in GC tissues and adjacent normal tissues by analyzing data from the GEO database, specifically the GSE63089 and GSE66229 datasets. Our analysis demonstrated a substantial upregulation of INHBA expression in GC tissues relative to adjacent normal tissues, as illustrated in Figure 1A. This finding was further corroborated by the significantly elevated expression of INHBA in stomach adenocarcinoma (STAD) tissues, as demonstrated through analysis using the GEPIA2 online tool (Figure 1B). Survival analysis conducted on GEPIA2 revealed that GC patients with high INHBA expression exhibited significantly poorer overall survival compared to those with low INHBA expression (Figure 1C). Given the related understanding of INHBA’s functional role in GC, we selected it as the target gene for further investigation. We verified INHBA mRNA expression in 34 pairs of GC tissues and their adjacent normal tissues using quantitative reverse transcription-polymerase chain reaction (qRT-PCR). The results revealed a statistically significant upregulation of INHBA mRNA in GC tissues compared to normal tissues (*P* < 0.05) (Figure 1D). To confirm these findings at the protein level, Immunohistochemistry (IHC) was performed on GC and adjacent normal tissues, demonstrating that INHBA protein was mainly localized in the cytoplasm of GC cells and significantly overexpressed in GC tissues (*P* < 0.001, *χ²* = 19.485) (Figure 1E, Table 1). Further we investigated the correlation between INHBA expression and the clinical characteristics of GC patients. Our analysis revealed that INHBA expression was independent of gender, age, tumor size, and tumor differentiation. However, it showed a significant association with tumor lesion, lymph node metastasis and TNM stage (Table 2). These findings suggest that INHBA overexpression is closely linked to a poor prognosis in GC patients, underscoring its potential as both a prognostic biomarker and a therapeutic target in GC. Finally we evaluated the mRNA and protein expression levels of INHBA in different GC cell lines and GES-1 cell using qRT-PCR and Western blot (WB) (Supplementary Figure 1A, Figure 1B).

**Figure 1.**
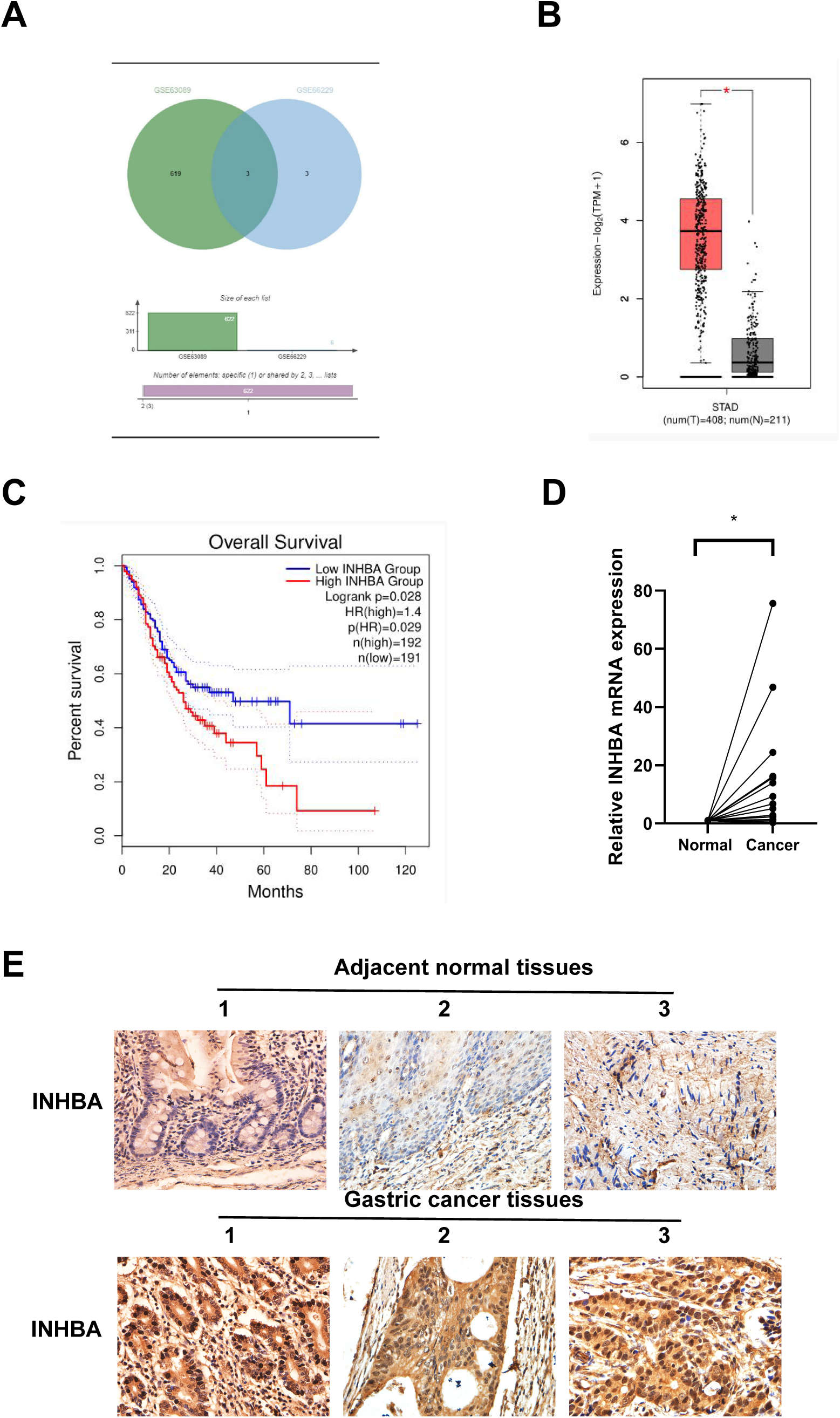
Increased INHBA expression is associated with poor prognosis in GC patients. (A) Expression of INHBA in the GEO database, including the GSE63089 and GSE66229 datasets. (B) INHBA is highly expressed in STAD tissues through the analysis of the online databases GEPIA2. (C) The survival analysis of INHBA expression levels and overall survival in patients with GC. (D) The expression levels of INHBA mRNA in 34 pairs of GC and adjacent normal tissue samples were detected by qRT-PCR. (E) Representative images showing INHBA protein expression levels in 50 pairs of GC and adjacent normal tissue samples, as detected by IHC. (magnification ×400).

**Table 1.** The expression level of INHBA protein in two groups.

| Groups | Cases | INHBA expression | | Positive<br>Rate(%) | $\chi^2$ | *P* |
| --- | --- | --- | --- | --- | --- | --- |
|  |  | Positive | Negative |  |  |  |
| Gastric cancer tissues | 50 | 38 | 12 | 76% | 19.485 | <0.001*** |
| Adjacent normal tissues | 50 | 16 | 34 | 32% |  |  |

**Table 2.** The relationship between the expression of INHBA in gastric cancer tissue and the clinicoapathological data of patients.

| Groups | Cases | INHBA expression | | $\chi^2$ | *P* |
| --- | --- | --- | --- | --- | --- |
|  |  | Positive | Negative |  |  |
| Gender |  |  |  |  |  |
| Male | 38 | 29 | 9 | 0.009 | 0.926 |
| Female | 12 | 9 | 3 |  |  |
| Age(years) |  |  |  |  |  |
| ≤60 | 8 | 6 | 2 | 0.005 | 0.942 |
| >60 | 42 | 32 | 10 |  |  |
| Tumor Lesion |  |  |  |  |  |
| Cardia | 8 | 2 | 6 | 13.581 | 0.000*** |
| Non-cardia | 42 | 36 | 6 |  |  |
| Tumor size |  |  |  |  |  |
| ≤4cm | 22 | 18 | 4 | 0.729 | 0.393 |
| >4cm | 28 | 20 | 8 |  |  |
| Differentiation |  |  |  |  |  |
| Poorly | 36 | 28 | 8 | 0.223 | 0.637 |
| Well | 14 | 10 | 4 |  |  |
| Lymph node metastasis |  |  |  |  |  |
| Absent | 11 | 5 | 6 | 7.214 | 0.007** |
| Present | 39 | 33 | 6 |  |  |
| TNM Staging |  |  |  |  |  |
| I | 1 | 0 | 1 | 10.147 | 0.017* |
| II | 13 | 7 | 6 |  |  |
| III | 26 | 21 | 5 |  |  |
| IV | 10 | 10 | 0 |  |  |

### Knockdown of INHBA suppresses the proliferation, migration, and invasion of GC cells and promotes apoptosis *in vitro*

To explore the functional role of INHBA in gastric cancer (GC), we first downregulated INHBA expression in NUGC-3 and Hs746T cell lines and assessed its impact on various cellular processes. Following transfection with siRNA targeting INHBA (siINHBA), a significant reduction in INHBA mRNA levels (Figure 2A, 2B) and protein levels (Figure 2C, 2D) was observed compared to the siRNA negative control (siNC). Among the four siRNA sequences of INHBA (si-281, si-382, si-499, si-822), we found that the effect of si-382 decline was the most obvious, so si-382 was selected for the subsequent knockdown of cell fuction experiments. The CCK-8 assay revealed that cell viability in NUGC-3 and Hs746T cells was significantly reduced following INHBA knockdown compared to the control group (Figure 2E, 2F). The wound healing assay demonstrated that the migratory capacity of INHBA knockdown cells was significantly impaired compared to the control group in both NUGC-3 and Hs746T cells after 48 hours (Figure 2G, 2H). Flow cytometry analysis revealed a significant increase in the number of apoptotic cells in both NUGC-3 and Hs746T cell lines following INHBA silencing compared to the control group (Figure 2I, 2J). Transwell assay results demonstrated a significant reduction in both migration and invasion capabilities in the INHBA knockdown group for both NUGC-3 and Hs746T cell lines (Figure 2K, L, M, N). These findings suggest that INHBA downregulation promotes apoptosis while inhibiting invasion, migration, and proliferation in GC cells.

**Figure 2.**
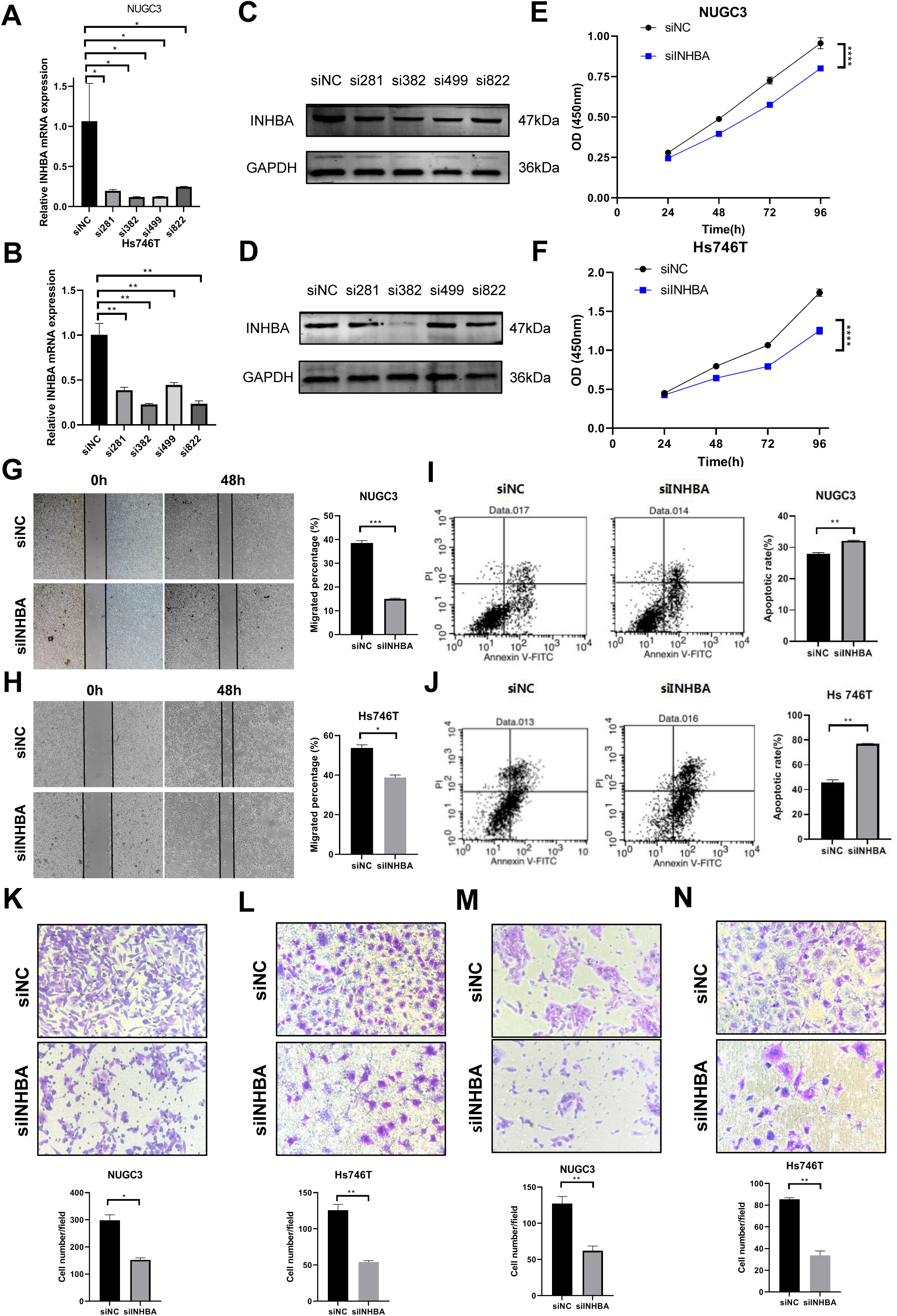
Knockdown of INHBA in GC cells suppresses cell proliferation, migration and invasion in vitro. (A, B) The expression of INHBA mRNA in NUGC-3 and Hs746T cells was detected by qRT-PCR. (C, D) The expression of INHAB protein in NUGC-3 and Hs746T was detected by WB. (E, F) The effect of INHBA knockdown on the proliferation of NUGC-3 and Hs746T cells was determined by the CCK-8 assay. (G, H) The effect of INHBA knockdown on the migration of NUGC-3 and Hs746T cells was assessed by the wound healing assay. Scale bar, 200 μm. (K, L) The effect of INHBA knockdown on the migration of NUGC-3 and Hs746T cells was evaluated using the Transwell migration assay. Scale bar, 100 μm. (M, N) The effect of INHBA knockdown on the invasion of NUGC-3 and Hs746T was assessed using Transwell invasion assay. Scale bar, 100 μm. (I, J) The effect of INHBA knockdown on the apoptosis of NUGC-3 and Hs746T cells was determined by flow cytometry. Data are presented as means ± SD. \**P* <0.05, \*\**P* <0.01, \*\*\**P* <0.001, \*\*\*\**P* <0.0001

### The over-expression of INHBA promotes the proliferation, migration, and invasion and suppresses the apoptosis of GC cells *in vitro*

To futher investigate the function of INHBA on cellular biological processes, we upregulated its expression in HGC-27 and AGS cell lines. Transfection with pcDNA3.1-INHBA significantly increased INHBA mRNA (Figure 3A, 3B) and protein levels (Figure 3C, 3D) compared to the pcDNA3.1-vector control. The CCK-8 assay demonstrated a significant increase in cell viability in HGC-27 and AGS cells following INHBA overexpression compared to the control group (Figure 3E, 3F). The wound healing assay revealed a significant enhancement in cell migration in HGC-27 and AGS cells following INHBA upregulation after 48 hours (Figure 3G, 3H). Flow cytometry analysis showed a significant increase in apoptotic cells in HGC-27 and AGS cells after INHBA upregulation (Figure 3I, 3J). Moreover, the Transwell assay demonstrated a significant increase in both migration and invasion in the INHBA-upregulated HGC-27 and AGS cells (Figures 3K, L, M, N). In summary, our findings suggest that INHBA acts as a pro-oncogene in GC, promoting invasion, migration, and proliferation, while inhibiting apoptosis.

**Figure 3.**
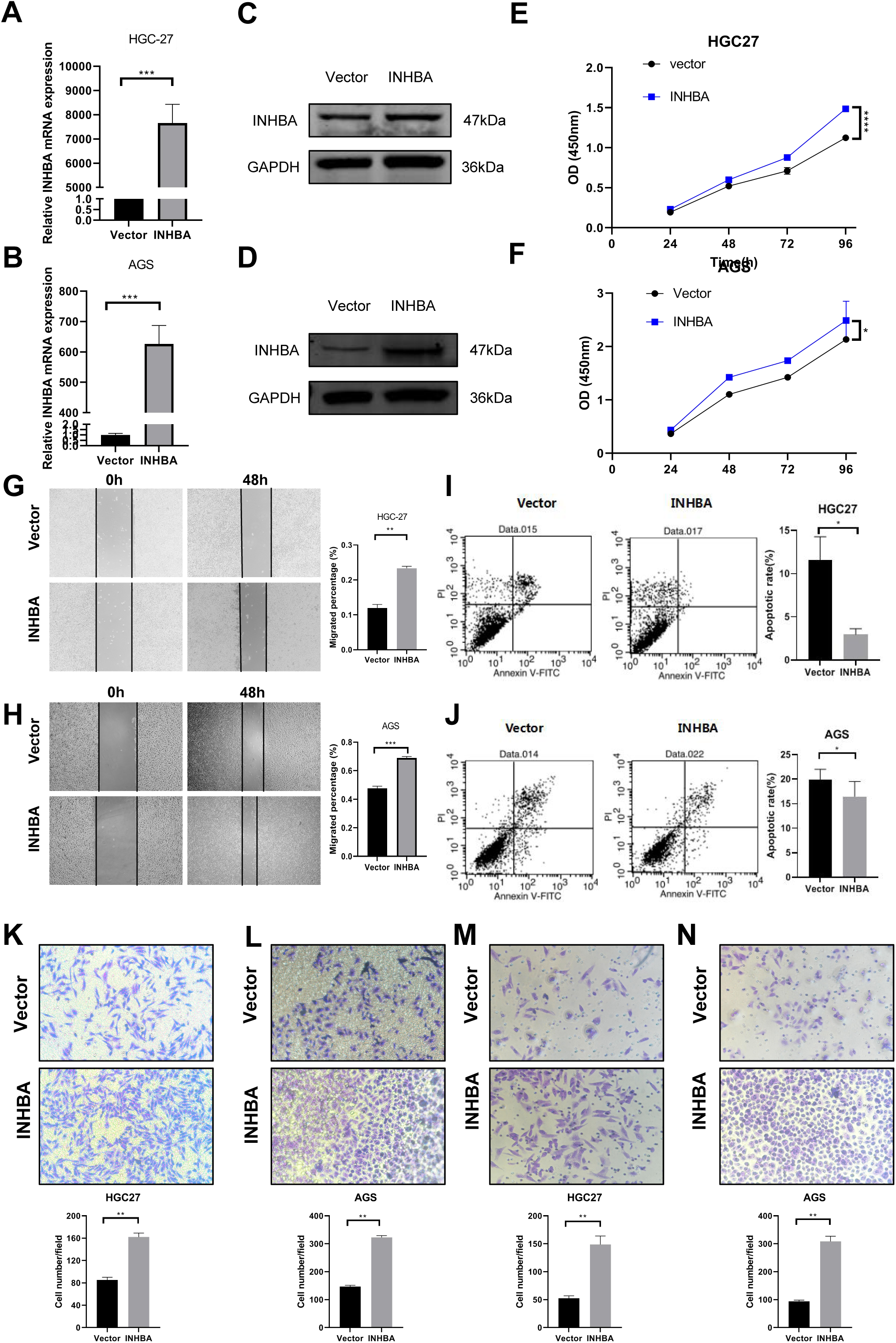
Overexpression of INHBA in GC cells and human gastric mucosa cells promotes cell proliferation, migration and invasion in vitro. (A, B) The expression of INHBA mRNA in HGC-27 and AGS was detected by qRT-PCR. (C, D) The expression of INHBA protein in HGC-27 and AGS was detected by WB. (E, F) The effect of INHBA overexpression on the proliferation of HGC-27 and AGS cells was assessed using the CCK-8 assay. (G, H) The effect of INHBA overexpression on the migration of HGC-27 and AGS cells was evaluated through a wound healing assay. Scale bar, 200 μm. (K, L) The effect of INHBA overexpression on the migration of HGC-27 and AGS cells was determined utilizing a Transwell migration assay. Scale bar, 100 μm. (M, N) The effect of INHBA overexpression on the invasion of HGC-27 and AGS cells was assessed via a Transwell invasion assay. Scale bar, 100 μm. (I, J) The effect of INHBA overexpression on the apoptosis of HGC-27 and AGS was analyzed by flow cytometry. Data are presented as means ± SD. \**P* <0.05, \*\**P* <0.01, \*\*\**P* <0.001, \*\*\*\**P* <0.0001

### INHBA over-expression promotes the growth of GC *in vivo*

To better investigate INHBA’s role *in vivo*, we established tumor xenografts in BALB/C nude mice using HGC-27 cells transfected with INHBA-overexpressing or control vectors. We first confirmed INHBA overexpression in the stably transfected cell lines by qRT-PCR and Western blot (Figure 1A-B), validating successful construction. HGC-27 cells stably transfected with INHBA overexpression or empty vectors were subcutaneously injected into BALB/C nude mice to generate tumor xenografts. Figures 4C-4F illustrated that INHBA-overexpressing tumors displayed a significantly higher growth rates, volumes, and weights compared to the vector group in xenograft models. Histopathological analysis of the xenograft tissues by HE staining demonstrated that the cellular architecture and morphological features closely resembled those observed in human GC tissues (Figure 4G). Immunohistochemical staining(IHC) for Ki67 revealed a significant increase in cell proliferative capacity in the INHBA-overexpression group compared to the vector group (Figure 4H). These findings demonstrated that INHBA overexpression promotes GC progression *in vivo*.

**Figure 4.**
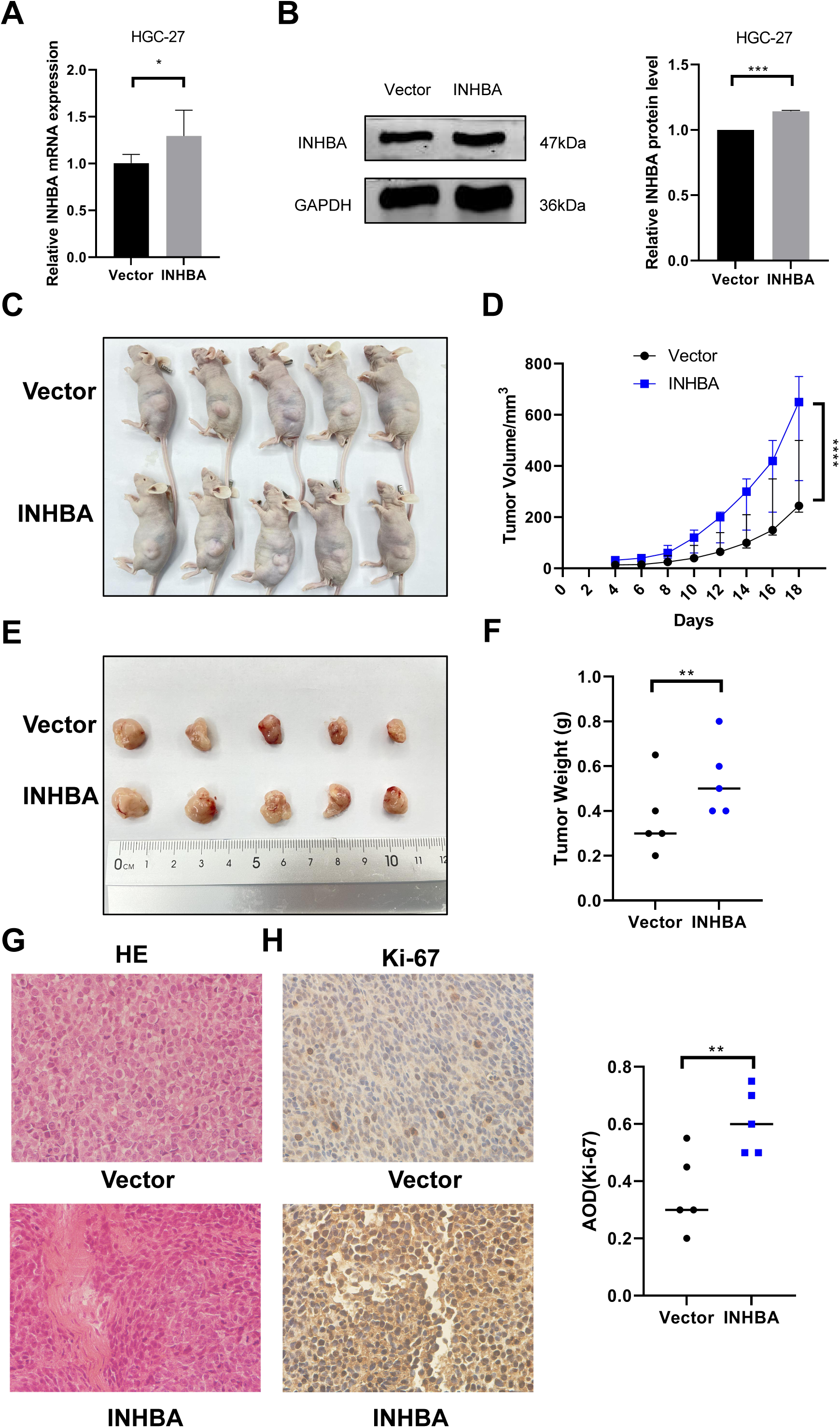
INHBA promotes the growth of GC *in vivo*. (A) Verification of stable bead over-expression efficiency was detected by qRT-PCR.(B) Verification of stable bead over-expression efficiency was detected by was detected by WB. (C) Nude mouse tumor-bearing model. (D) Tumor growth curve. (E, F) Images (E) and weight (F) of xenografted tumors were analyzed. (G) Representative images of tumor tissue sectionsstained with HE. (magnification ×400). (H) Representative image of Ki67 IHC staining in tumor tissue sections. (magnification ×400). Data are shown as means ± SD.\**P* <0.05, \*\**P* <0.01, \*\*\**P* <0.001, \*\*\*\**P*<0.0001

### ITGA6 is the potential target of INHBA in GC

To clarify the pathophysiological role of INHBA in GC, we conducted transcriptome RNA sequencing (RNA-seq) on NUGC-3 cells transfected with either non-targeting siRNA (siNC) or INHBA-targeting siRNA (siINHBA). RNA-seq results revealed INHBA silencing elicited widespread transcriptome dysregulation, with 1,958 differentially expressed genes identified, comprising 1,036 up-regulated and 922 down-regulated genes, as demonstrated in the gene expression heatmap and volcano plot (Figure 5A, B). Through the analysis of RNA-seq-identified downregulated genes (log2Fold Change>1, *P*<0.05), along with the 3,742 genes that are highly expressed in GC within the GEPIA2 database, 10 genes were identified as potential candidates for binding to the INHBA protein(Figure 5C). Since INHBA is predominantly localized extracellularly, we selected the membrane proteins from candidate genes as potential targets for INHBA regulation. ITGA6 was identified as the only membrane protein and exhibited broad distribution across multiple cellular compartments in GC cells among the 10 candidate genes. Bioinformatic analysis using the GEPIA2 database demonstrated constitutive overexpression of ITGA6 in gastric adenocarcinoma specimens (Figure 5D), with multicompartment localization patterns showing predominant distribution in cytomembrane, cytoplasm, and nucleus domains (Figure 5E). Therefore, we speculate that ITGA6 is a potential downstream target gene. Co-IF and Co-IP assays were then conducted to demonstrate the interaction between INHBA and ITGA6. The Immunofluorescence co-localization analysis revealed that INHBA and ITGA6 co-localize and are mainly found in the cytoplasm and cytomembrane of GC cells (Figure 5F). Co-IP assay confirmed direct INHBA-ITGA6 interaction in GC cells (Figure 5G). To confirm INHBA-mediated regulation of ITGA6 expression, we performed bidirectional INHBA modulation (knockdown and overexpression) in GC cells. INHBA knockdown decreased ITGA6 mRNA and protein in NUGC-3 cells (Figures 5H, I). Conversely, both ITGA6 mRNA and protein expression levels were increased after INHBA expression was upregulated in AGS cells (Figures 5J, K). These results suggest that INHBA regulates downstream gene expression at both the transcriptional and translation levels. In summary, these results revealed an functional interaction between INHBA and ITGA6 in GC, and that ITGA6 may act as a potential modification target of INHBA in GC.

**Figure 5.**
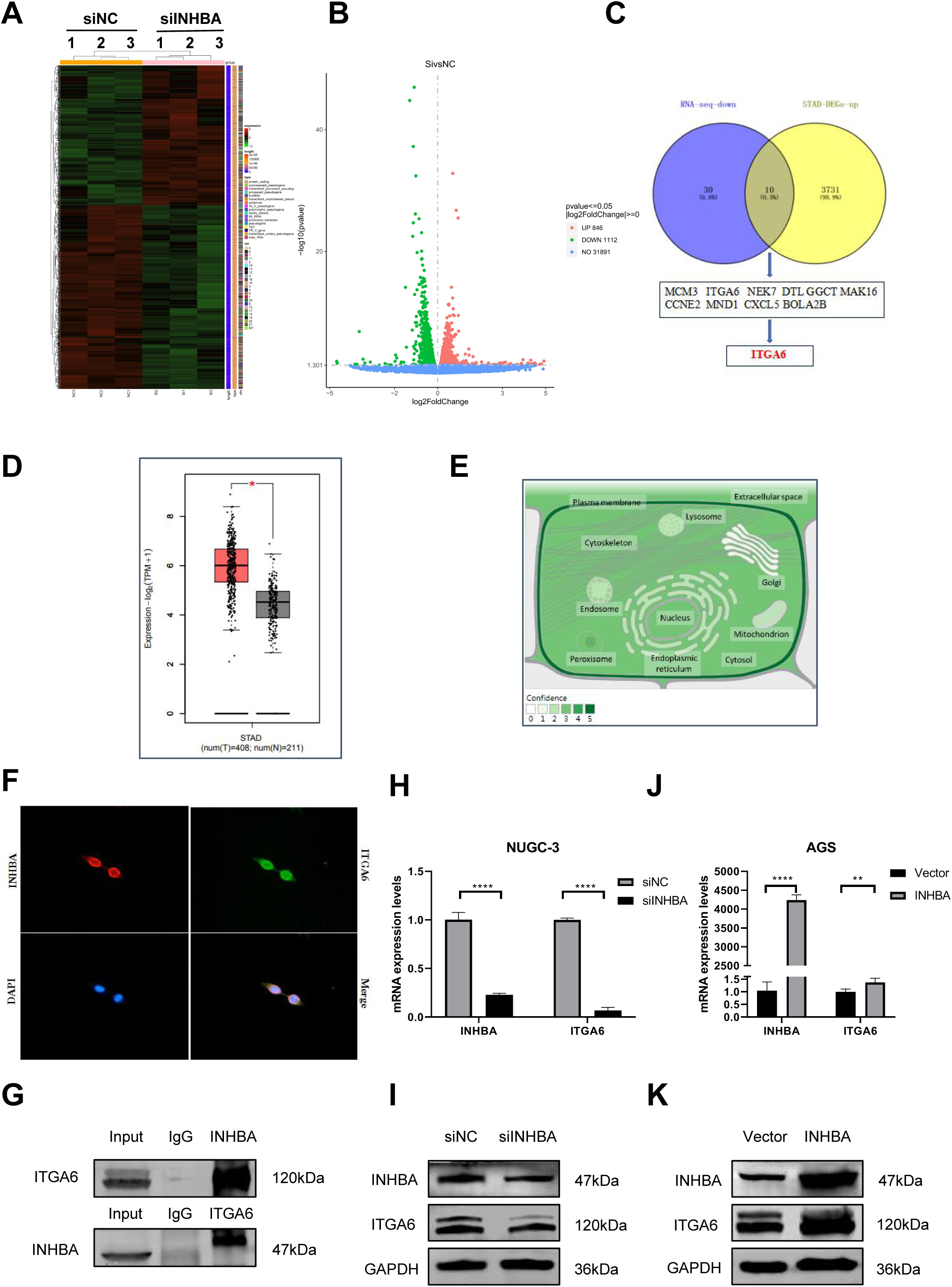
ITGA6 is the regulation target of INHBA. (A) Differential expression genes (DEGs) heat map identified by RNA-seq. (B) The number of DEGs identified by RNA-seq. (C) Overlap analysis of RNA-seq and STAD-DEGs-up identified genes. (D) Expression ITGA6 in GC paired tissue cohort from the GEPIA2 database. (E) Subcellular localization of ITGA6 from GeneCards. (F, G) the interaction between INHBA and ITGA6 was verified by Co-IF (F) and Co-IP (G) experiments. Scale bar, 10μm. (H) qRT-PCR was used to detect ITGA6 mRNA expression after INHBA knockdown in NUGC-3 cells. (I) WB was used to detect ITGA6 protein expression after INHBA knockdown in NUGC-3 cells. (J) qRT-PCR was used to detect ITGA6 mRNA expression after INHBA overexpression in AGS cells. (K) WB was used to detect ITGA6 protein expression after INHBA overexpression in AGS cells. Data are presented as means ± SD. \**P* <0.05, \*\**P* <0.01, \*\*\**P* <0.001, \*\*\*\**P*<0.0001

### ITGA6 is highly expressed in GC and interacts with INHBA

Next, we evaluated ITGA6 expression in both GC cell lines and clinical specimens. Quantitative analysis revealed that majority of GC cell lines exhibited significantly elevated ITGA6 mRNA expression compared to the normal gastric mucosa cells (Figure 6A). Western blot analysis identified NUGC-3 as exhibiting the highest ITGA6 protein expression among all tested cell lines (Figure 6B). IHC analysis of 30 paired clinical specimens demonstrated elevated ITGA6 expression in GC tissues versus adjacent normal tissues (Figure 6C). These results suggest ITGA6 is highly expressed in GC tissues and various GC cell lines. Further studies should illustrate how INHBA-ITGA6 interactions contributes to gastric carcinogenesis. Cell lines were selected based on baseline ITGA6/INHBA expression level. Following functional screening of four ITGA6-targeting siRNAs in AGS cells with the highest ITGA6 expression, si-ITGA6-614 demonstrated superior silencing efficiency and was prioritized for further analyses (Figure 6D-E). We then conducted reciprocal modulation strategies in GC cell models: NUGC-3 received ITGA6 overexpression (pcDNA3.1-ITGA6) and INHBA knockdown (siRNA-INHBA), while AGS underwent ITGA6 silencing (siRNA-ITGA6) and INHBA overexpression (pcDNA3.1-INHBA). qRT-PCR and Western blot were used to identify the variations in INHBA expression in each group. The results demonstrated ITGA6 overexpression could counteract INHBA knockdown effect in NUGC-3 cells (Figure 6F-G). Similarly, ITGA6 silencing can also mitigate INHBA overexpression effect in AGS cells (Figure 6H-I). These results reveal that ITGA6 can influence INHBA expression in GC cells.

**Figure 6.**
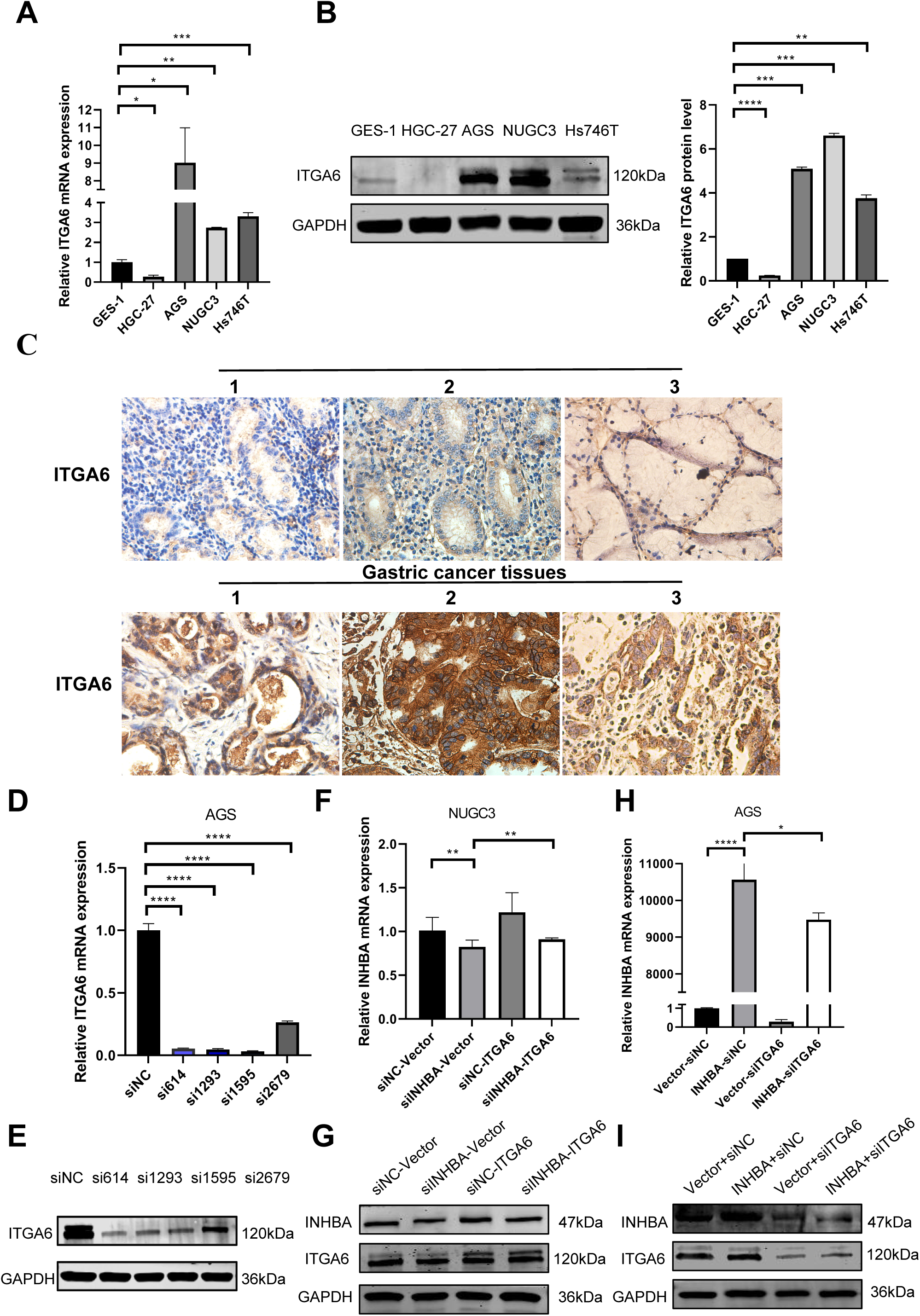
Expression of ITGA6 and its interaction with INHBA in GC. (A) The basal expression of ITGA6 mRNA in GC cell lines and GES-1 was detected by qRT-PCR. (B) The basal expression of ITGA6 protein in cell lines was detected by WB. (C) The expression of ITGA6 protein in 30 pairs of GC paired tissues was detected by IHC. (magnification ×400). (D-E) Four siRNA-ITGA6 sequences were transfected simultaneously into AGS cells. (F-G) The relative expression level of INHBA mRNA and protein were detected by qRT-PCR and WB after instantaneous co-transfection of INHBA siRNA and ITGA6 overexpression plasmid in NUGC-3. (H-I) The relative expression level of INHBA mRNA and protein were detected by qRT-PCR and WB after instantaneous co-transfection of ITGA6 siRNA and INHBA overexpression plasmid in AGS. Data are presented as means ± SD. \**P* <0.05, \*\**P* <0.01, \*\*\**P* <0.001, \*\*\*\**P*<0.0001

### INHBA targets ITGA6 to activate the MAPK signaling pathway in GC promotion

To futher explore the molecular mechanism of INHBA and ITGA6, we selected the top 1000 genes co-expressed with INHBA and ITGA6 in GC from TCGA database and analyzed the signaling pathways controlled by them. KEGG pathway enrichment analysis based on the Database for Annotation, Visualization and integrated Discovery (DAVID) shown that gene clusters were significantly enriched in MAPK signaling pathway (Figure 7A). Therefore, we speculated that INHBA activates ITGA6 via MAPK signaling pathway to drive gastric cancer progression. We then examined ITGA6’s ability to rescue INHBA knockdown-induced MAPK signaling pathway suppression. Western blot analysis revealed concomitant downregulation of MAPK cascade components (p-MEK, p-ERK1/2) following INHBA silencing in NUGC-3 cells. At the same time, ITGA6 overexpression rescued INHBA knockdown-mediated suppression of MAPK phosphoproteins such as p-MEK and p-ERK1/2 (Figure 7B). These results demonstrated that INHBA regulates the activation of the MAPK signaling pathway via ITGA6. Functional interaction studies revealed ITGA6 modulate INHBA-induced oncogenic phenotypes. Functional rescue assays showed ITGA6 overexpression rescued INHBA silencing-induced attenuation of cell proliferation, migration, and invasion in NUGC-3 cells (Figure 7C-G). Similarly, the down-regulation of ITGA6 expression can reduce the enhancement of proliferation, migration and invasion of AGS cells caused by INHBA overexpression (Figure 7H-L). In conclusion, The INHBA/ITGA6 axis drives gastric carcinogenesis through MAPK signaling pathway activation, with ITGA6 serving as the indispensable mediator of INHBA’s oncogenic activity. In summary, it was found that elevated INHBA can promote the expression of the ITGA6 mRNA and protein, which futhermore activate the MAPK signaling pathway and eventually contribute to the progression of GC.

**Figure 7.**
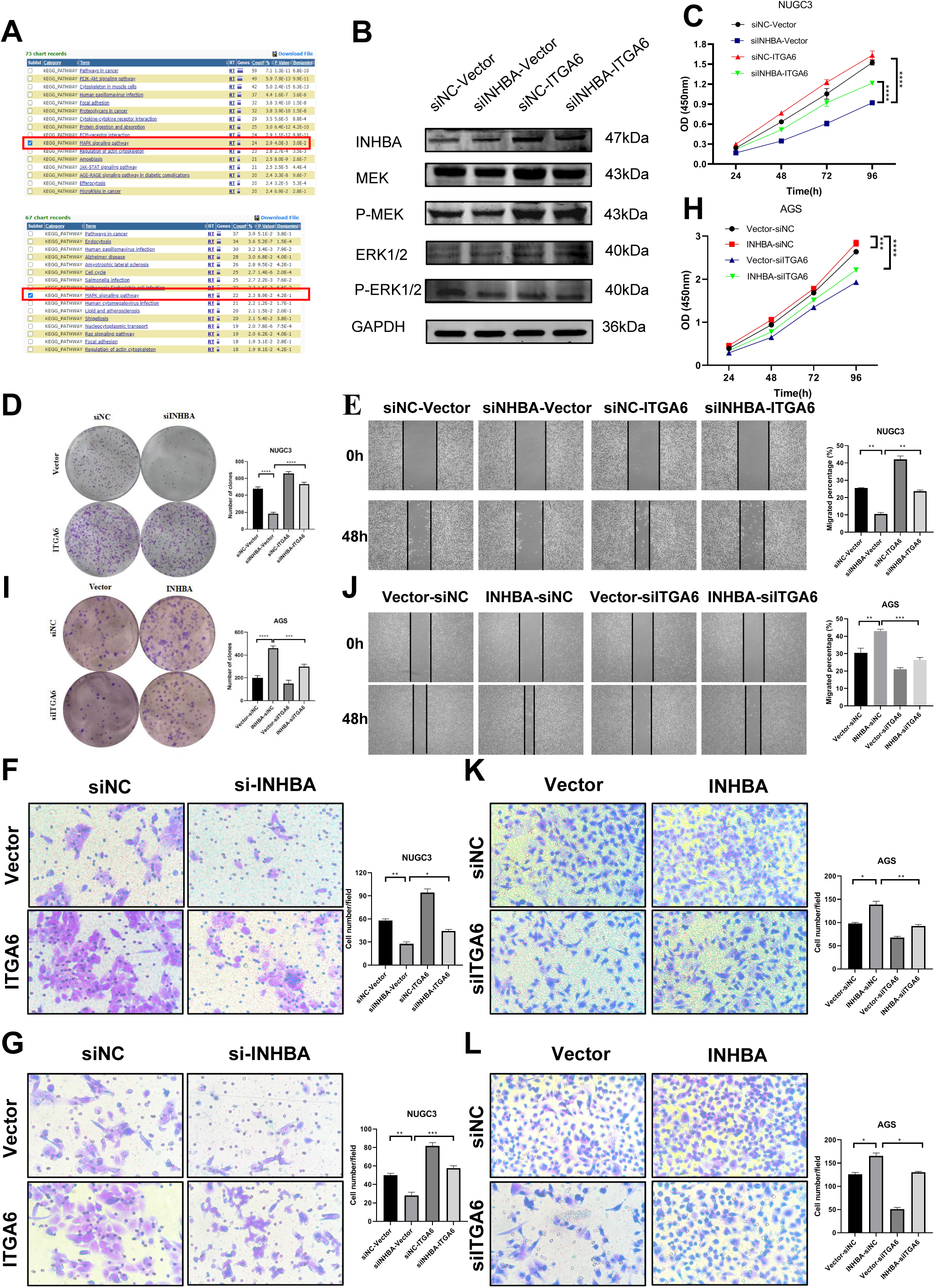
Oncogenic function of INHBA depends on ITGA6 and MAPK signaling pathway. (A) KEGG enrichment result of the differentially expressed gene sets from the GEPIA2 database. (B) The expression levels of MAPK signaling pathway related proteins were detected by WB after instantaneous co-transfection of INHBA siRNA and ITGA6 overexpression plasmid in NUGC-3. (C, D, E, F, G) The effects of ITGA6 on INHBA in NUGC-3 cells were detected by CCK-8 assay (C), colony formationassay (D), wound healing assay (E), Scale bar, 200 μm, Transwell migration assay (F) and Transwell invasion assay (G). Scale bar, 100 μm. (H, I, J, K, L) The effects of ITGA6 on INHBA in AGS cells were detected by CCK-8 assay (H), colony ormation assay (I), wound healing assay J), Scale bar, 200 μm, Transwell migration assay (K) and Transwell invasion assay (L). Scale bar, 100 μm.. Data are presented as means ± SD. *\*P <0.05, **P <0.01, ***P <0.001, ****P <0.0001*

## Discussion

This study shows that INHBA overexpression in gastric cancer patients correlates with poor prognosis. Functional validation through gain-of-function assays demonstrated INHBA contributed to tumorigenic phenotypes *in vitro* and *in vivo*. INHBA knockdown suppresses GC cell proliferation, migration, invasion and promotes apoptosis, contrasting with the oncogenic effects of its overexpression. Mechanistically, INHBA drives gastric carcinogenesis by upregulating ITGA6, activating MAPK signaling pathway to promote tumor progression.

Previous studies showed significantly higher INHBA expression in GC tissues compared to adjacent normal tissues. These results were significantly related to lymph node metastasis and TNM staging[19]. In this study, clinical and pathological characteristics of 50 pairs GC and adjacent normal tissues were assessed, revealing results in agreement with previous reports. Importantly, We found that the positive expression rate of INHBA was significantly lower in cardiac cancer compared to non-cardiac cancer, according to our study results. The clinical significance of this finding in decision-making treatment approaches for cardiac and non-cardiac cancers need futher in-depth and comprehensive research. In conclusion, We proposed that INHBA may be a clinically useful, prognostic indicator of poor patient survival.

To confirm the functional influence of INHBA on gastric cancer cellular processes, experimental data revealed that genetic silencing of INHBA markedly inhibited GC cell proliferation, migration, and invasion while increasing apoptosis. In contrast, overexpression of INHBA resulted in conversely biological outcomes. Furthermore, animal studies utilizing immunodeficient nude mice demonstrated that overexpression of INHBA drove the progression of gastric tumors. Former evidence indicated that INHBA overexpression enhances malignant behaviors of GC cells in vitro models, whereas genetic inhibition of INHBA suppresses these oncogenic phenotypes[15,18]. Several studies have shown that INHBA are closely associated with various types of cancer, such as esophageal squamous cancer [5], colorectal cancer [20], breast cancer [21], cervical cancer [22] and ovarian cancer [23]. Consistent with our data, these results support INHBA’s role in promoting tumorigenic properties (proliferation, metastasis) during gastric carcinogenesis.

Following this, we attempted to discover the genes targeted by INHBA in order to uncover its biological mechanisms in GC pathogenesis. Integrated analysis of downregulated genes in RNA-seq data, STAD-DEGs-up overlap, database screening and literature veiwing identified ITGA6 as a candidate gene. Western blot analysis following co-immunoprecipitation and Co-immunofluorescence experiments demonstrated physical interaction between INHBA and ITGA6 in GC cells, a novel discovery in the field. As a member of the integrin family, ITGA6, also known as α6-integrin, CD49f, is a cell surface protein that mediates cell-to-cell and cell-to-stroma adhesion, which is vital to cell proliferation, migration, survival, and differentiation[24]. Previous studies have demonstrated that ITGA6 is highly expressed in various cancers. Moreover, the crucial role of ITGA6 has been verified in ovarian cancer, bladder cancer, and pancreatic cancer [25–27]. Notably, these data aligned with our ITGA6 observations in gastric cancer as shown in Figure 6A-C. To investigate how INHBA modulates ITGA6 in gastric cancer, we revealed that INHBA governs ITGA6 expression through both transcriptional and translational mechanisms as shown in Figure 5H-K. In summary, INHBA potentially drives gastric cancer progression by enhancing ITGA6 expression through mRNA and protein binding.

KEGG pathway enrichment analysis based on the Database for Annotation, Visualization and integrated Discovery (DAVID) showed that INHBA and ITGA6 co-expressed gene clusters were significantly enriched in MAPK signaling pathway. The MAPK signaling pathway acts as a pivotal mediator in relaying extracellular cues to intracellular effectors, with its cascade dynamics frequently dysregulated through epigenetic modifications in diverse pathological contexts, most notably during oncogenic transformation. Studies have demonstrated that MAPK pathway is vital to the process of gastric cancer development, including cellular proliferation, invasion, migration, and metastasis [28]. Previous studies have revealed that MAPK signaling pathway-related proteins are widely distributed in various types of cancer, such as colorectal cancer [29], esophageal squamous cancer [30], pancreatic cancer [31] and hepatocellular carcinoma[32]. MEK(Mitogen activated protein kinase) is a key kinase in MAPK signaling pathway, which can be activated by upstream kinases and then phosphorylated and activated downstream ERK(Extracellular regulated protein kinase) to regulate cell growth, differentiation and proliferation [32]. Therefore, we hypothesized that INHBA could activate MAPK signaling pathway to promote the progression of GC. We discovered that INHBA knockdown-mediated suppression of MAPK pathway activation was reversed by ITGA6 overexpression, which itself directly stimulated MAPK signaling pathway. Rescue studies demonstrated that the INHBA/ITGA6/MAPK cascade critically governs GC cell malignant behaviors, including proliferation, migration, and invasion. These findings demonstrate that the INHBA/ITGA6/MAPK axis drives gastric cancer progression, offering diagnostic and therapeutic targets and underscoring INHBA’s broader oncogenic role.

However, this study has some limitations requiring future investigation. Although mechanistically confined to MAPK signaling pathway, our findings do not preclude the potential engagement of other pathways in INHBA-mediated oncogenic processes. The specific mechanisms and interactions of these signaling pathways need to be further explored. While this study establishes the therapeutic potential of INHBA/MAPK/ITGA6 axis inhibition in gastric cancer, pharmacological validation using clinical-grade inhibitors needs pre-clinical efficacy assessments *in vitro* and *in vivo* models.

In conclusion, this investigation demonstrates INHBA’s pivotal oncogenic function in driving gastric cancer cell malignancy (proliferation, migration, and invasion), pioneering in elucidating its molecular pathogenesis via ITGA6 interaction-mediated MAPK signaling pathway. This study identifies INHBA as a potential diagnostic and prognostic biomarker. Moreover, the INHBA/ITGA6 axis affects the development of GC by regulating the MAPK signaling pathway, emerging as a druggable target, providing molecular rationale theoretical basis for GC precision therapy.

## Acknowledgments

This work was supported by Hebei Provincial Government-funded Provincial Medical Excellent Talent Project (ZF2023025, ZF2024134, ZF2025045, ZF2025048, and ZF2025051), Hebei Natural Science Foundation (H2022206292, H2024206140), Key R&D Program of Hebei Province (223777103D and 223777113D), Hebei Province County General Hospital Appropriate Health Technology Promotion Project (20220018) and other projects of Hebei Province (SGH201501) and Medical Science Foundation of Hebei University (2024B03).

---

**Supplementary Figure 1.**
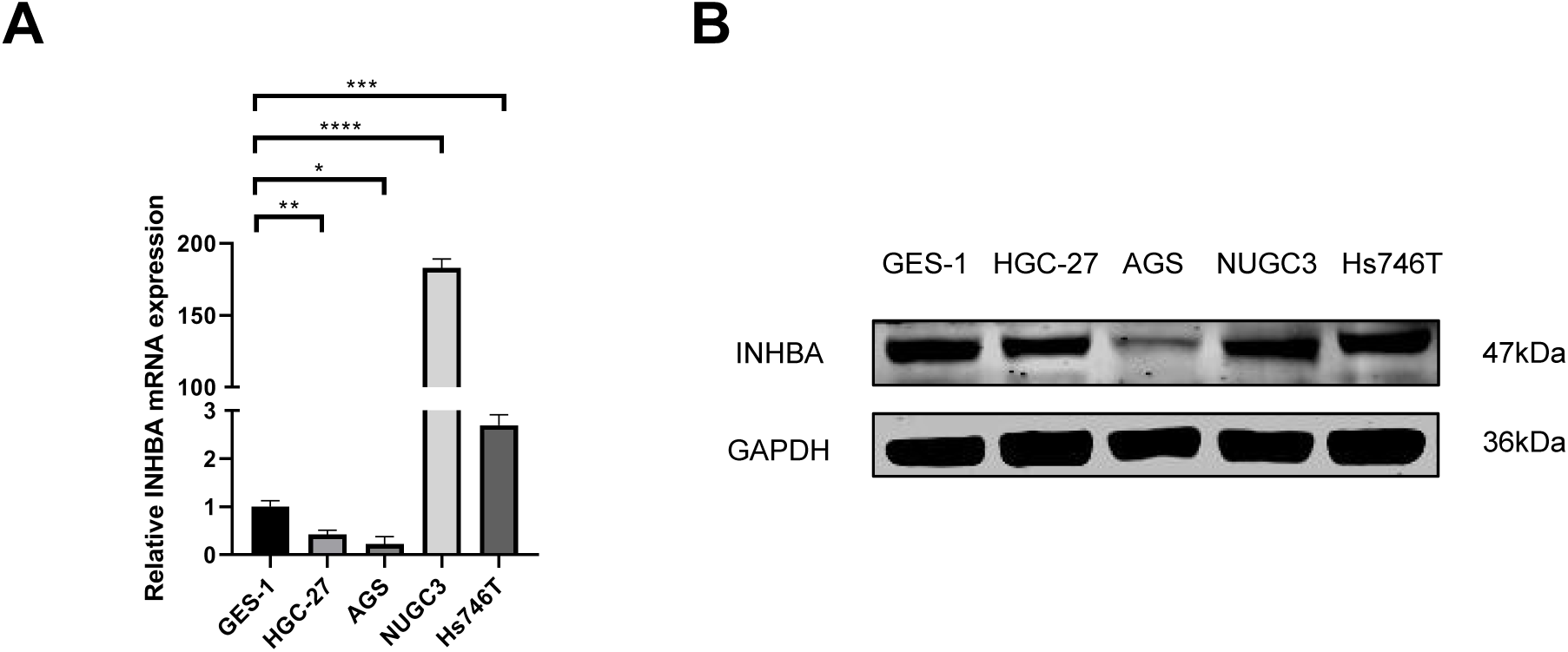
(A) The expression of INHBA mRNA in GC cell lines and GES-1 was detected by qRT-PCR. (B) The basal expression of INHBA protein in GC cell lines and GES-1 cell lines was detected by WB. Data are presented as means ± SD. \**P* <0.05, \*\**P* <0.01, \*\*\**P* <0.001, \*\*\*\**P*<0.0001

**Supplementary Table 1.**
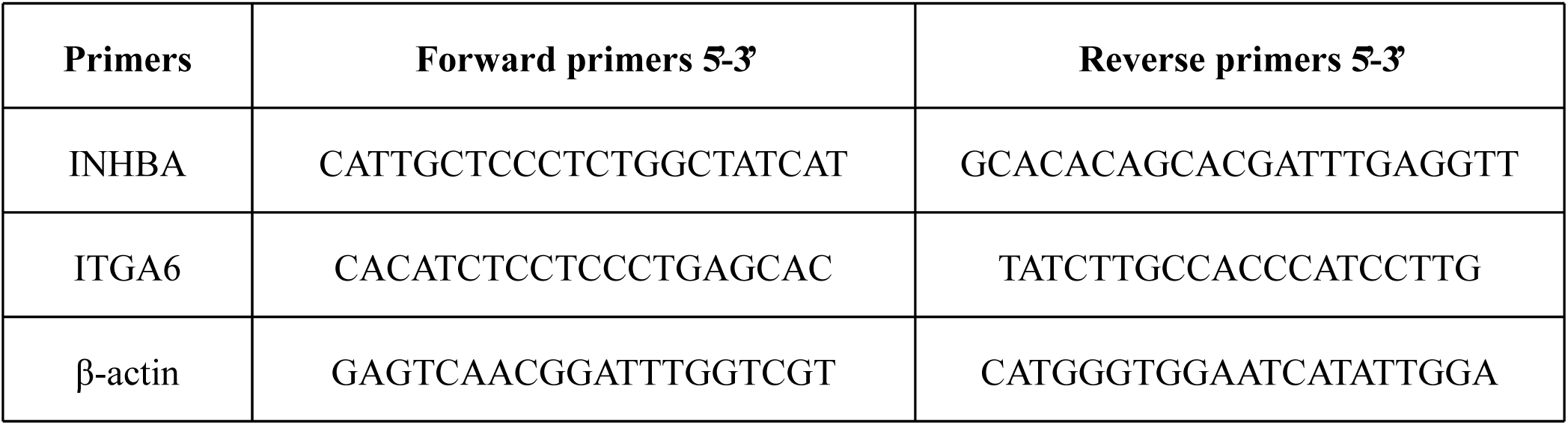
The primer sequences of INHBA, ITGA6, β-actin.

